# The expectant brain – being pregnant causes changes in brain morphology in the early postpartum period

**DOI:** 10.1101/2021.06.29.450283

**Authors:** Natalia Chechko, Jürgen Dukart, Svetlana Tchaikovski, Christian Enzensberger, Irene Neuner, Susanne Stickel

## Abstract

There is growing evidence that pregnancy may have a significant impact on the maternal brain, causing changes in its structure. However, the patterns of these changes have not yet been systematically investigated. Using voxel-based (VBM) and surface-based morphometry (SBM), we compared a group of healthy primiparous women (n = 40) with healthy multiparous mothers (n = 37) as well as nulliparous women (n = 40). Compared to the nulliparous women, the young mothers showed decreases in gray matter volume in the bilateral hippocampus/amygdala, the orbitofrontal cortex/subgenual prefrontal area, the right superior temporal gyrus, the right insula, and the cerebellum. However, these pregnancy-related changes in brain structure did not predict the quality of mother-infant attachment at either 3 or 12 weeks postpartum, nor were they more pronounced among the multiparous women. SBM analyses showed significant cortical thinning especially in the frontal and parietal cortices, with the parietal cortical thinning likely potentiated by multiple pregnancies. We conclude, therefore, that the widespread morphological changes seen in the brain shortly after childbirth reflect substantial neuroplasticity. Also, the experience of pregnancy alone may not be the underlying cause of the adaptations for mothering and caregiving. As regards the exact biological function of the changes in brain morphology as well as the long-term effect of pregnancy on the maternal brain, further longitudinal research with larger cohorts will be needed to draw any definitive conclusions.

**Significance Statement:** Biological adaptations during pregnancy affect the maternal brain. Here, we evaluated morphological changes in the brain of mentally healthy young mothers within the first four days of childbirth. Compared to the nulliparous women, the young primiparous and multiparous mothers demonstrated a substantial reduction in gray matter volume in brain areas related to socio-cognitive and emotional processes. Cortical alterations due to pregnancy-related adaptations are not the underlying cause of mother-infant attachment.

## Introduction

Pregnancy is characterized by a remarkable increase in circulating steroid hormones, including estradiol, progesterone and cortisol (1, 2). Both in animals and humans, these continually increasing hormones are thought to change the pregnant brain by crossing the blood-brain barrier (1, 3). In female mice, for instance, gestation and lactation periods have been seen to induce strong, albeit transient, gray matter concentration hypertrophy within key regions involved in emotion, motivation, reward, and mnestic functions (4). There are no longitudinal studies in humans involving continuous observation of changes from prior to conception, through pregnancy and the postpartum period. However, at least two independent studies have concluded that pregnancy is accompanied by significant decreases in brain size (5) and gray matter volume (GMV) (6). A number of studies have focused on brain structure changes only during the postpartum period, not assessing brain morphology either before or during pregnancy. In one study, the increase in GMV was observed between 2-4 weeks and 3-4 months postpartum (7). According to another study, a postpartum increase in GMV is already visible between the acute (within 1-2 days postpartum) and subacute postpartum (4–6 weeks after childbirth) periods (8), suggesting that neuroplasticity processes start very early in the postpartum period. Collectively, these studies suggest that multiple areas (the (para)-hippocampus, the fusiform gyrus (6), the temporal lobe (6, 7), the amygdala (7), the precuneus/posterior cingulate (6, 8), the medial prefrontal/orbitofrontal areas (6, 7), the inferior frontal gyrus (6–8), the pre- and postcentral gyri (7, 8), the insula (6, 7), the hypothalamus (7), the striatum and thalamus (7, 8)) are involved in peripartum/postpartum adaptation. Thus, a network comprising the regions involved in emotion regulation (e.g. prefrontal/anterior cingulate cortices, amygdala, hippocampus), sensory and social information processing (e.g. superior temporal sulcus and gyrus, precuneus/posterior cingulate cortex), reward (e.g. striatum, insula, hypothalamus, amygdala), memory and learning (e.g. prefrontal cortex, amygdala, hippocampus) as well as stress processing and regulation (e.g. prefrontal/anterior cingulate cortices, amygdala, hippocampus) is suggested to underlie human maternal brain plasticity (9).

However, the exact biological function of the brain regions associated with pregnancy and postpartum changes is not clear. In line with the findings in animal models (4), two studies in humans suggest (6, 7) that these changes are associated with motherhood, facilitating the development of maternal attachment to the child. In particular, the bilateral insula, the inferior frontal gyrus, the striatum, the superior temporal gyrus, the thalamus and the amygdala are considered to be active in the so-called maternal brain (10). However, while these changes are thought to be adaptive, preparing new mothers for their new role, their possible contribution to the development of mental disorders cannot be ruled out as the changes in brain structure and the development of postpartum psychiatric disorders co-occur in time (11). Importantly, some regions underlying human maternal brain plasticity (9) are also thought to play a role in the development of postpartum depression (PPD) with PPD-related functional abnormalities having been reported in the amygdala, the insula, and the orbitofrontal and dorsomedial prefrontal cortices (for reviews, see 3, 12). Understanding the biological functions of pregnancy- and postpartum period-associated alterations in brain structure, therefore, is of great significance, as they can be linked to both advantages and risks for new mothers. However, there are a number of issues that need a closer examination. First, we do not know exactly how pregnancy and the postpartum period change the maternal human brain. Studies addressing structural neuroplasticity in pregnancy and the postpartum period do not report, likely due to their own methodological problems, the exact areas of overlap. Given that the postnatal adaptation processes involving structural change begin very early in the postpartum period (5, 8) and last up to 6 months (5) or longer (6), the investigation time points are likely to influence the results. And these studies mostly included groups that were very heterogeneous in terms of the time points at which they were investigated. For instance, Hoekzema et al. (6) examined women between 1 and 4 months postpartum, disregarding the fact that the maternal brain 1 month postpartum may differ from the one 4 months postpartum. In addition, the sample sizes of these studies were not very high, typically including between 9 and 19 participants (5, 7, 8). Also, the very early postpartum period (1-2 days postpartum) was assessed in only one study (8, 13). Assessing the very early postpartum period (e.g. the first few days postpartum) is of particular relevance as it may shed valuable light on the entire spectrum of changes associated with pregnancy. This period is also crucial for the development of mother-infant attachment, facilitated by increased responsivity toward the infant’s cues (14–19). The first few days postpartum are also associated with some negative feelings or mood swings (the so-called baby blues). Lasting up to 4 to 6 weeks after childbirth, the subacute postpartum period puts a women at risk for PPD (DSM 5, 20). Whether pregnancy-induced alterations in plasticity and brain morphology have long-lasting effects, extending beyond the reproductive event itself (21), is yet to be determined. Further research is needed to draw definitive conclusions in this regard and, also, to find out whether multiple pregnancies exert long-lasting, pregnancy-related effects on the brain. None of the previous studies compared first-time pregnant women with those with multiple pregnancies. Nor were there comparisons with control groups comprising yet-to-be-pregnant women. According to some studies (6, 7), changes in a woman’s brain structure across pregnancy are linked to certain aspects of maternal caregiving, such as the absence of hostility toward the child (6) or the mother’s subjective perception of parenting and the baby (7). Hoekzema et al. (6) assessed maternal attachment by means of the Maternal Postnatal Attachment Scale (MPAS; 22) in relationship to the pregnancy-related GMV changes, although MPAS was employed only once and that too retrospectively, for the first six months of motherhood (6). As maternal attachment develops over time and can be influenced by a number of factors, a single assessment of attachment with respect to changes in the brain structure following pregnancy may not be sufficiently reliable.

In view of all this, the overarching goal of this project was to explore pregnancy-related GMV changes in young mothers (n = 78), within the very first few days postpartum, compared to a group of nulliparous women (n = 40). The postpartum mothers underwent magnetic resonance imaging (MRI) measurements at 2 days postpartum on average. To determine whether the number of pregnancies can have any impact on the specific pregnancy-related changes in brain volume, we compared primiparous women with their multiparous counterparts (2 to 3 children). Given that the data were collected as part of a longitudinal study, including observation of maternal attachment at several time points within the first 12 weeks postpartum, we also focused on the link between pregnancy-related changes and the development of attachment based on multiple MPAS assessments. Factors such as baby blues, duration of pregnancy, birth mode and child’s gender were also taken into account. Between the young mothers and controls, we expected to find differences in several regions underlying functional plasticity of the maternal brain, including the hippocampus, and the prefrontal and temporal cortices. We hypothesized these differences to be more pronounced in multiparous mother. In addition, we expected an association between the maternal brain and the development of attachment toward the infant over the 12-week postpartum period.

## Results

### Behavioral results

Table 1 summarizes the characteristics of the subdivided group according to parity, including age, gestational age, birth mode, experience of baby blues, EPDS scores and MPAS scores.

**Table 1.** Sample characteristics of postpartum women.

| | Primiparous<br>($n = 41$) | Multiparous<br>($n = 37$) | Statistical test |
| --- | --- | --- | --- |
|  | *Mean (SD)* | *Mean (SD)* |  |
| Age | 31.24 (4.09) | 32.57 (4.66) | $t(76) = -1.33, p = .189$ |
| Gestational age in days | 276.68 (10.90) | 275.43 (10.63) | $t(76) = 5.12, p = .610$ |
| Child's birth weight | 3281.10 (454.91) | 3431.08 (508.45) | $t(76) = -1.36, p = .173$ |
| MPAS 3 weeks pp | 86.07 (4.69) | 84.05 (5.82) | $t(76) = 1.69, p = .094$ |
| Quality of attachment | 40.45 (2.68) | 39.11 (3.41) | $t(76) = 1.93, p = .058$ |
| Absence of hostility | 22.85 (2.14) | 22.76 (1.89) | $t(76) = 0.20, p = .841$ |
| Pleasure in interaction | 23.05 (1.72) | 22.19 (2.89) | $t(57.76) = 1.57, p = .122$ |
| MPAS 6 weeks pp | 86.20 (4.33) | 83.73 (6.31) | $t(62.84) = 1.99, p = .051$ |
| Quality of attachment | 40.35 (31.10) | 39.49 (3.45) | $t(76) = 1.56, p = .251$ |
| Absence of hostility | 22.80 (1.92) | 22.05 (2.31) | $t(76) = 1.54, p = .127$ |
| Pleasure in interaction | 23.05 (1.41) | 22.19 (2.85) | $t(51.71) = 1.65, p = .104$ |
| MPAS 9 weeks pp | 85.93 (4.79) | 83.65 (6.29) | $t(66.98) = 1.78, p = .079$ |
| Quality of attachment | 40.53 (3.43) | 39.22 (3.59) | $t(76) = 1.64, p = .106$ |
| Absence of hostility | 22.55 (2.14) | 22.16 (2.48) | $t(76) = 0.74, p = .463$ |
| Pleasure in interaction | 22.95 (1.68) | 22.30 (2.60) | $t(60.75) = 1.29, p = .200$ |
| MPAS 12 weeks pp | 86.22 (4.66) | 85.05 (6.19) | $t(76) = 0.95, p = .348$ |
| Quality of attachment | 40.55 (3.34) | 40.11 (3.51) | $t(76) = 0.57, p = .573$ |
| Absence of hostility | 22.80 (2.42) | 22.27 (2.38) | $t(76) = 0.97, p = .336$ |
| Pleasure in interaction | 23.25 (1.31) | 22.43 (2.89) | $t(49.51) = 1.58, p = .121$ |
| EPDS at birth | 3.39 (2.48) | 4.32 (2.45) | $t(76) = -.167, p = .099$ |
| EPDS 3 weeks pp | 4.34 (3.09) | 4.27 (2.28) | $t(73.18) = 0.16, p = .908$ |
| EPDS 6 weeks pp | 3.51 (2.44) | 3.70 (2.59) | $t(76) = -0.33, p = .739$ |
| EPDS 9 weeks pp | 2.71 (2.39) | 3.27 (2.68) | $t(76) = -0.98, p = .330$ |
| EPDS 12 weeks pp | 2.24 (1.88) | 2.65 (2.29) | $t(76) = -0.86, p = .394$ |
|  | *Percent* | *Percent* |  |
| Birth mode | | | Fisher's exact = 5.34, $p = .125$ |
| Spontaneous | 68.3 | 78.4 |  |
| Ventouse | 9.8 | - |  |
| C-section | 12.2 | 18.9 |  |
| Emergency C-section | 9.8 | 2.7 |  |
| Experience of baby blues (yes/no) | 48.8 | 54.1 | $\chi^2(1) = .216, p = .642$ |

Note. SD: Standard deviation; MPAS: Maternal Postnatal Attachment Scale (22); EPDS: Edinburgh Postnatal Depression Scale (23); pp: postpartum.

Primiparous and multiparous participants did not differ significantly in age, gestational age, birth mode and the experience of baby blues (yes/no, based on self-report and MBQ cut-off score > 10) (see Table 1). The groups did not differ in the self-reported values of mother-child attachment total scores and the respective subscales, although there was a trend for primiparous mothers to have slightly higher MPAS scores than multiparous mothers 6 weeks postpartum. In addition, the groups did not differ in the self-reported depression scores at any time point.

Repeated measures ANOVA was used within each group to analyze the postpartum course of the total mother-child attachment scores as well as the three subscales at all four time points. There was no significant effect of time in either group regarding the total scores (primiparous: F(2.41, 96.3) = 0.98, p = .935; multiparous: F(3, 108) = 2.00, p = .118). Additionally, no significant interaction was detected between the subscales and the time points (primiparous: F(6, 234) = 0.26, p = .955; multiparous: F(4.25, 152.91) = 1.95, p = .101).

Repeated measures ANOVA showed a significant effect of time regarding the EPDS values in both groups (primiparous: F(4, 160) = 7.28, p < .001, η_p_^2^ = .154; multiparous: F(4, 144) = 4.03, p = .004, η_p_^2^ = .101). Within the primiparous group, Bonferroni-corrected pairwise comparisons revealed significantly higher depressive values at 3 weeks compared to 9 (p = .003, d =.59) and 12 weeks postpartum (p < .001, d =.820), as well as higher values at 6 weeks compared to 12 weeks (p = .024, d =.583). Within the multiparous group, depressive values at 12 weeks were significantly lower compared to after childbirth (p = .018, d = .705), 3 weeks (p = .01, d = .709) and 6 weeks postpartum (p = .041, d = 0.234).

### Whole-brain VBM results

#### GMV alterations in primiparous participants

We tested whether pregnancy and childbirth lead to changes in the brain structure. In a whole-brain analysis, using t-contrast from random-effects GLM, we found a symmetrical pattern of significant GMV decrease in three clusters in the primiparous compared to the nulliparous women (Figure 1A, Table S1). The first cluster (20607 voxel) included regions of the cerebellum extending to the inferior temporal gyrus, the fusiform gyrus and the inferior occipital gyrus. The second cluster (12159 voxel) included the basal ganglia, the subgenual and orbitofrontal cortices, the anterior and midcingulate cortices, the thalamus, the insula, the temporal cortex, the hippocampus, the parahippocampal gyrus and the amygdala. Further decreases were observed in a cluster (3988 voxel) predominantly located in the superior parietal lobule, the paracentral and postcentral gyri, the midcingulate cortex, the precuneus and the cuneus. No significant GMV increase was found in the primiparous compared to the nulliparous women.

**Figure 1.**
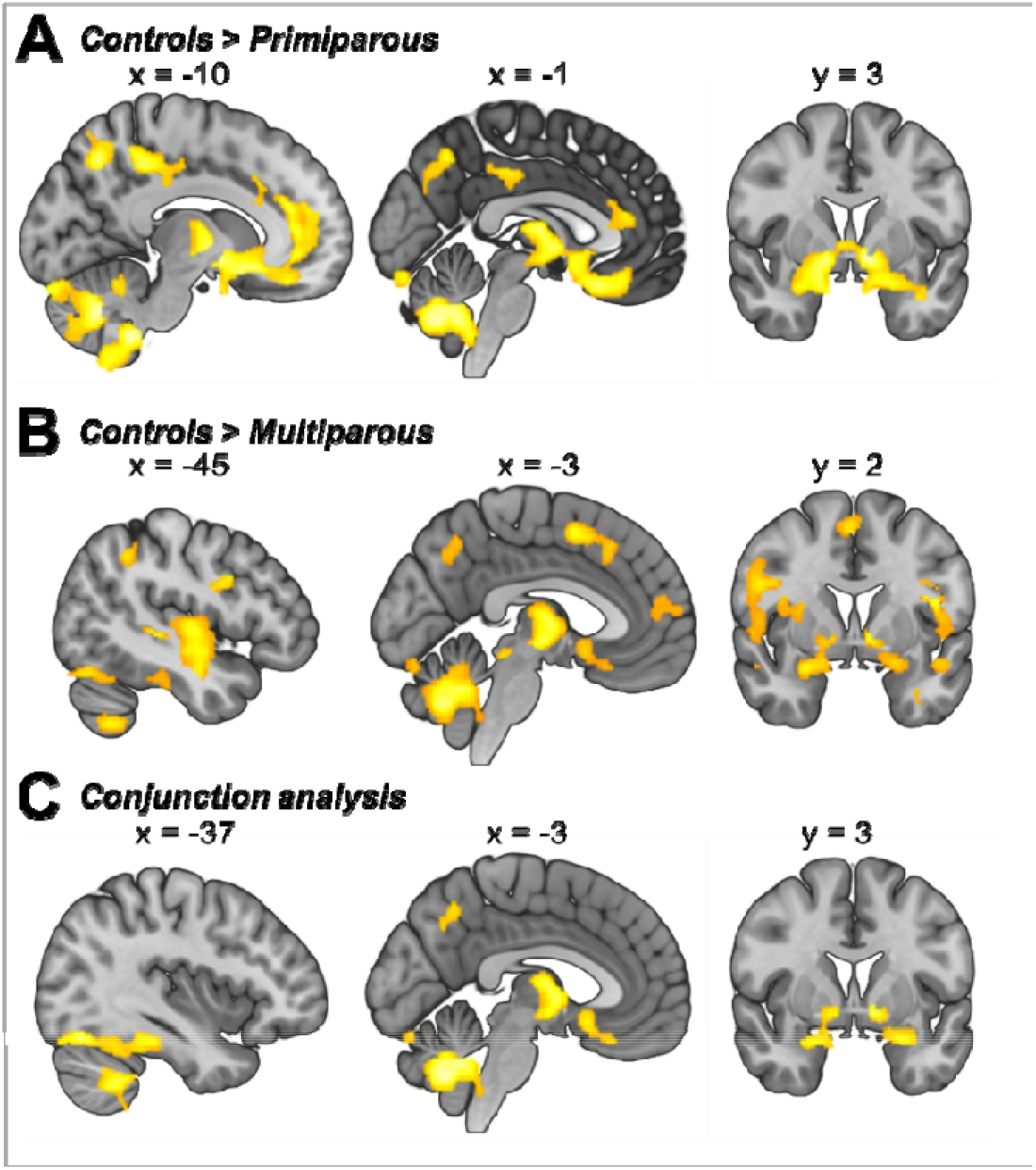
Gray matter volume differences in control subjects compared to A) primiparous and B) multiparous women. C) Brain regions linked to pregnancy based on a conjunction analysis (controls > primiparous ⍰ controls > multiparous). Illustration of clusters emerging from t-contrast from random-effects GLM, p < .05, cluster-level FWE correction k = 920)

#### GMV alterations in multiparous participants

Seeking to determine if GMV changes are potentiated by multiple pregnancies, we compared the nulliparous to the multiparous women. The whole-brain analysis of GMV differences between control subjects and multiparous women revealed a symmetrical pattern of significant decrease in multiparous women in six clusters (Figure 1B, Table S2). The first cluster (36157 voxels) encompassed widespread GMV differences located in the cerebellum and the temporal lobe with extensions observed in the fusiform gyrus, the basal ganglia, the amygdala, the hippocampus and the parahippocampal gyrus as well as the insula, the inferior frontal gyrus and the orbital cortex. In both hemispheres of the multiparous women, significant decreases were seen in one cluster (7842 voxel) encompassing the temporal cortex, the rolandic operculum, the insula, the thalamus, the basal ganglia, the rectal gyrus, the parahippocampal gyrus and the hippocampus, in two clusters (4838 and 1972 voxel) including the dorsal frontal, orbital and cingulate cortices, and in two clusters (4300 and 1379 voxel) encompassing the parietal and occipital cortices. An additional cluster (3520 voxels) in the right hemisphere included the rolandic operculum, the insula, the inferior frontal gyrus (p. triangularis) and the putamen. No significant GMV increase was found in the multiparous compared to the nulliparous women.

### GMV alterations in primiparous compared to multiparous participants

Based on the GMV changes potentiated by multiple pregnancies, we further tested for long-lasting GMV changes by comparing primiparous and multiparous women. No significant group differences were identified between primiparous and multiparous participants in the t-contrast from random-effects GLM with a cluster-forming threshold at voxel-level p < .001. Likewise, when applying a more liberal cluster-forming threshold at voxel-level p < .005 as well as p < .01, no significant differences were found in the pairwise comparisons.

### GMV alterations linked to pregnancy

To isolate the GMV difference-based links to pregnancy in general, we performed a conjunction analysis across both contrasts (control > primiparous ⍰ control > multiparous) and found the pregnancy-related GMV differences to be most pronounced in a cluster (16736 voxel) encompassing the cerebellum, the bilateral fusiform gyrus and the temporal cortex extending to the parahippocampal gyrus. Another cluster (4003 voxel) was found in both hemispheres in the thalamus, the temporal pole, the pallidum and the rectal gyrus extending to the basal ganglia, the thalamus, the amygdala, the hippocampus, the parahippocampal gyrus, the insula and the medial orbital gyrus. The third cluster (1507) included the left hemispheric superior parietal gyrus and the inferior gyrus (Figure 1C, Table S3).

### Effects of baby blues and hormonal contraception on GMV differences

To determine whether the experience of baby blues is associated with GMV changes, we first compared all postpartum women who experienced symptoms of baby blues (n = 40) with those who had not (n = 38) (and vice versa) and found no significant main effect of baby blues on GMV. The comparison between primiparous women with baby blues (n = 20) and those without (n = 21) did not yield significant results either, nor did the comparison between multiparous women with baby blues (n = 20) and those without (n = 17).

In addition, to investigate whether ovarian hormones per se can modify GMVs, we compared control subjects who used hormonal contraception (n = 19) with those who did not (n = 22) and found no significant GMV differences between these groups.

### Effect of child’s gender on GMV

Forty-one of the postpartum women carried a girl (25 primiparous women), and 37 a boy (16 primiparous). Considering the previously observed effects of fetal sex on cognitive changes in pregnant women (24), we additionally compared the brain structures of the mothers of boys with those of the mothers of girls. We found no significant effect of gender on GMV and no interaction between parity group and gender.

### Multivariate regression analyses

To determine whether maternal behavior toward the child is linked to structural changes due to pregnancy and childbirth, we examined the GMV in relation to maternal attachment. Kernel ridge regression was used to test whether the MPAS total score (Figure 2) and its subscales (quality of attachment, absence of hostility, and pleasure in interaction; Table 5) at 3 and 12 weeks postpartum were predicted by whole-brain GMV at birth in the whole postpartum sample. The analysis showed that neither the overall quality of mother-infant attachment nor the subscales were predicted by GMV at any time point.

**Table 2.** Prediction of quality of attachment, absence of hostility and pleasure in interaction at 3 and 12 weeks postpartum by gray matter volume at birth.

|  | Quality of attachment | Absence of hostility | Pleasure in interaction |
| --- | --- | --- | --- |
| 3 weeks postpartum | $r = .03, p = .369$ | $r = -.01, p = .712$ | $r = .18, p = .07$ |
| 12 weeks postpartum | $r = .02, p = .389$ | $r = -.07, p = .653$ | $r = .16, p = .116$ |

**Figure 2.**
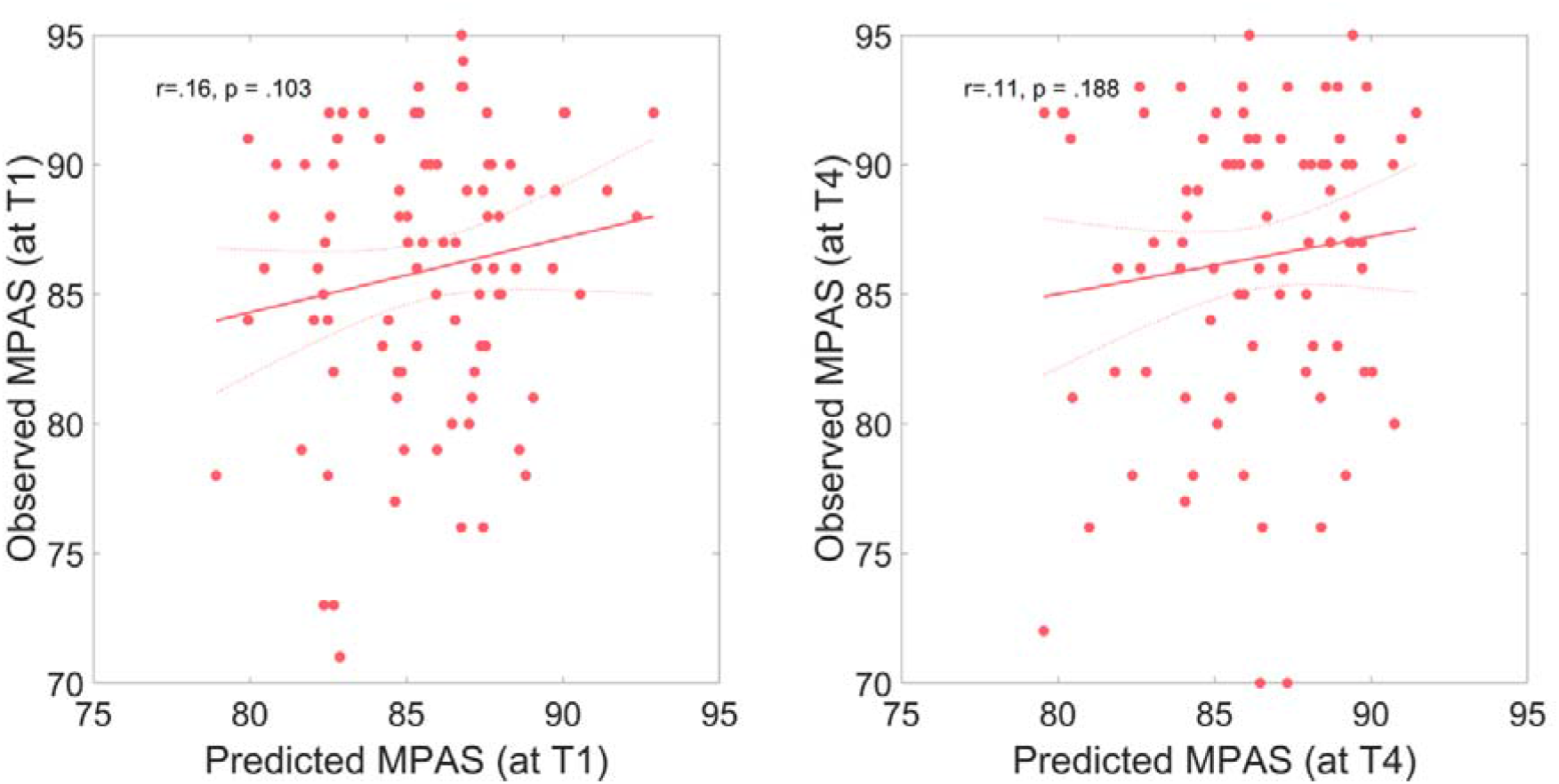
Multivariate prediction of Maternal Postpartum Attachment Scale (MPAS) scores based on the gray matter volume at birth. Predicted versus actual scores are plotted at A) 3 weeks postpartum (pp) and B) 12 weeks pp. The solid red line indicates a linear fit and the dashed lines the 95% confidence interval.

### Results of whole-brain SBM

#### Differences in cortical thickness between control subjects and primiparous women

Finally, we assessed the structural changes in the maternal brain by means of surface-based analyses. In a whole-brain analysis, using t-contrast from random-effects GLM, we found reduced cortical thickness in the bilateral dorsolateral prefrontal cortex, the bilateral superior parietal gyrus as well as the right inferior parietal sulcus and the supramarginal gyrus in the primiparous compared to the nulliparous women (Figure 3A, Table S4).

**Figure 3.**
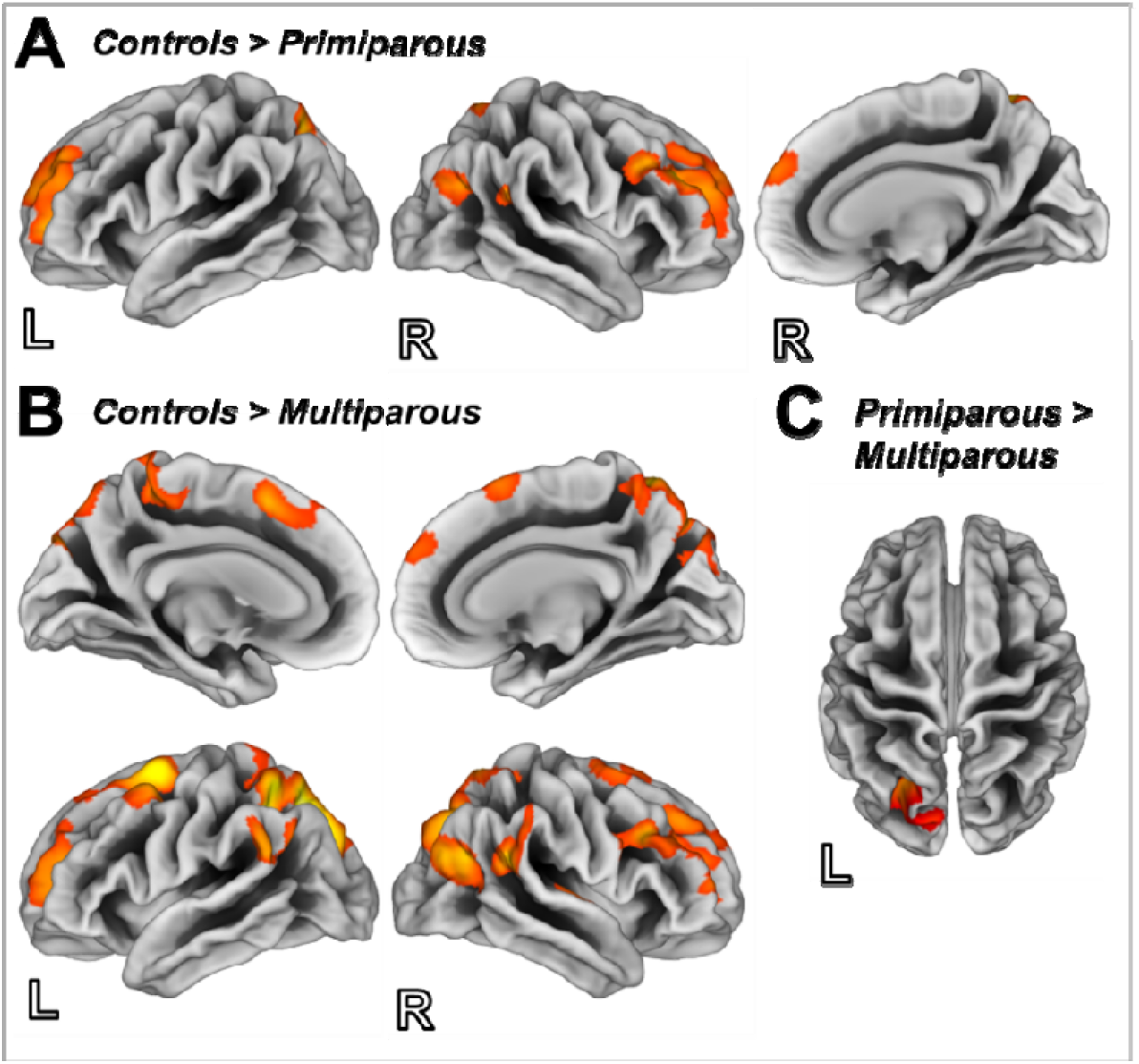
Surface maps depicting differences in cortical thickness in control subjects compared to A) primiparous women and B) multiparous women, and C) in primiparous compared to multiparous women. Illustration of clusters emerging from t-contrast from random-effects GLM, p < .05, cluster-level FWE correction, k = 116 voxels).

#### Differences in cortical thickness between control subjects and multiparous women

Multiple pregnancies were found to result in reduced cortical thickness in the bilateral dorsolateral as well as the dorsomedial prefrontal cortices. In addition, both the parietal and temporal cortices were reduced in the multiparous compared to the nulliparous women (Figure 3B, Table S4).

#### Differences in cortical thickness between primiparous and multiparous women

Investigating the possible long-lasting changes in brain structure, we found multiparous women to exhibit less cortical thickness, compared to their primiparous counterparts, in the superior parietal gyrus and the inferior parietal sulcus (Figure 3C, Table S4).

## Discussion

During pregnancy and the early postpartum period, mothers undergo psychological and biological adaptations that trigger processes involving brain plasticity. In order to assess the structure underlying maternal brain plasticity in relation to pregnancy, we compared the brain structure of postpartum women within the very early postpartum period with that of nulliparous women. Shortly after childbirth (on overage 2 days postpartum), young mothers displayed an extensive GMV decrease in the bilateral hippocampus/amygdala, the orbitofrontal cortex/the subgenual area (BA 25), the bilateral temporal lobe, the insula, the basal ganglia and the cerebellum. In addition, the surface-based analyses revealed significant cortical thinning in the frontal and parietal cortices, with the parietal cortical thinning likely potentiated by multiple pregnancies.

The brain areas affected by pregnancies are not random, as they (the medial prefrontal cortex and the temporal lobe in particular) have been shown to be related to socio-cognitive processes and emotion perception (25) and be part (insula, amygdala, inferior frontal gyrus, superior temporal gyrus) of the neuroanatomy of the so-called maternal brain (10) and the reward system (basal ganglia). Pregnancy-induced neural plasticity occurs both in humans and other mammals, affecting similar functions. For instance, in female mice, gestation and lactation periods have been found (similarly to our findings in humans) to alter the structure of brain regions involved in the regulation of social behaviors and social reward (medial preoptic area) as well as those involved in emotions (amygdala), motivation and reward (caudate nucleus, orbitofrontal cortex) and mnestic functions (hippocampus) (4). Both in humans and other mammals, the sex steroid hormones and corticosteroids are suggested to be the main mediators of the observed morphometric changes (1, 3). In particular, the amygdala, the hippocampus and the prefrontal cortex are densely covered with receptor cells for corticosteroid and ovarian hormones (26–28). As these brain areas are associated with a variety of major functions in humans, the effects of pregnancy on emotional and cognitive processes, and also on the development of maternal behavior, are likely to be significant, although they have yet to be sufficiently investigated. In addition, as any phase of the brain in transition (for reviews, see 3, 12), pregnancy increases the risk of psychiatric disorders (3). In this context, a link between pregnancy-related GMV changes and postpartum psychiatric disorder cannot be completely ruled out given their co-occurrence in time.

### Effects of pregnancy-associated brain alterations on emotional and cognitive functions and the development of PPD

The effects of pregnancy on emotional and cognitive demands are supported by several studies (e.g. 29, 30). A meta-analysis by Davies et al. (31) reports, for instance, that the overall cognitive function (memory, executive function and attention) is poorer in pregnant women compared to non-pregnant women, especially during the third trimester. This is hardly surprising given that our results along with those of animal studies suggest that pregnancy triggers (at least transitorily) a reduction of hippocampal volume (4, 32, 33). That being said, the reports pertaining to the overall cognitive function in pregnant and postpartum women are not always consistent (e.g. 34, 35). For definitive conclusions in this regard, longitudinal studies are needed to simultaneously examine brain structure and cognitive function during pregnancy and the postpartum period. As regards the effect of pregnancy on emotion processing and stress regulation, the subgenual anterior cingulate (in addition to the amygdala and the hippocampus) is related to the regulation of emotions and stress (36). In line with that, a recent study by our group found postpartum women with higher cumulative hair cortisol concentration in the last trimester of pregnancy to have lower activation in the subgenual ACC during emotional interference involving anxious emotions shortly after childbirth (30). In addition, the subgenual anterior cingulate cortex, the hippocampus and the amygdala are known to be related to affective disorders (37). For instance, the reduction in the mean subgenual anterior cingulate cortex gray matter volume in patients with bipolar and major depression disorders, compared to healthy controls, is well documented (38, 39) and thought to be linked to a reduction in glia cells (40). This raises the question as to whether these morphological changes seen shortly after childbirth are related (particularly in women with an increased risk of affective disorders) to the development of conditions like PPD. We sought to address this question in the RiPoD study (Schnakenberg et al., 2021) but, based on the multimodal neuroimaging data obtained shortly after childbirth, did not find any structural and functional brain differences between participants who would develop PPD and healthy controls. However, the number of PPD cases investigated in the RiPoD study thus far is relatively small (n = 21) and, therefore, more research is needed to gain insight into the matter. Finally, given that we did not find any GMV differences between primiparous and multiparous women, we cannot conclude that morphological changes in the early postpartum period are more pronounced in multiparous mothers. Nevertheless, multiparous women, compared to their primiparous counterparts, showed more pronounced cortical thinning particularly in the parietal cortices. Changes in cortical thickness, with age-related thinning in particular, are observed in normal development and aging (41). Rapid thinning, however, is often apparent in middle-aged groups (42) with diseases such as Alzheimer’s (e.g. 43), and also in individuals with familial risk for depression (44) and emerging or manifest depression disorder (45, 46).

### Is mother’s behavior related to structural changes in the brain during pregnancy?

According to some previous studies, the typical behavior of a new mother is associated with structural changes in the maternal brain during and immediately after pregnancy (6, 7). For instance, the brain volume reduction seen in humans at 1-4 months postpartum is thought to be predicted by the retrospectively reported absence of hostility in new mothers in the first 6 months following childbirth (6), suggesting a link between pregnancy-related GMV reduction and the quality of attachment to the child. Choosing a different approach, we sought to determine if the Maternal Postpartum Attachment Scale (MPAS) scores at 3 or 12 weeks postpartum could be predicted based on the gray matter volume at childbirth. However, we did not find any significant association. A crucial difference between our study and the one by Hoekzema et al. (6) is that, in their study, the examination of the mother’s brain took place considerably later (1-4 months postpartum) than in ours (1-2 days postpartum). Although Hoekzema et al. attributed the GMV alterations to pregnancy-related changes, we are of the opinion that what they actually examined was the postpartum brain. We argue also that the experience of pregnancy may not be the sole underlying cause of adaptations for mothering and caregiving. Pregnancy likely changes the brain in such a way that the link between the brain and the felt attachment evolves during the process of becoming a mother and is probably contributed to, in no small measure, by the interaction with the child. Given that fathers also develop attachment and caregiving skills (47, 48), other neurobiological mechanisms, and not only pregnancy-related brain structure alterations, are likely responsible for the transition to effective parenting. For instance, in a study by Kim et al. (49), biological fathers showed GMV increases in the hypothalamus, the amygdala, the striatum, and the lateral prefrontal cortex between 2-4 weeks and 12-16 weeks after childbirth. Thus, the development of attachment is likely to be a time-dependent process rather than being driven by pregnancy-related changes in brain structure.

### Limitations

In sum, MRI examinations of women within a very narrow time frame after childbirth showed the early postpartum period to be associated with pronounced alterations in brain structure. These alterations did not predict the development of attachment to the child in the first few weeks after delivery, nor did they appear to be potentiated by multiple pregnancies. However, the ways in which the networks and the brain structures restore themselves following childbirth cannot be determined by our data given our study’s lack of longitudinal MRI assessment and the lack of pre-pregnancy MRI, which is a major limitation of the current work. In addition, the MPAS was obtained for the first time 3 weeks postpartum (thus, 2-3 weeks after the neuroimaging data were collected). Additional neuroimaging data at 3 or 12 weeks postpartum would be needed to determine if postpartum alterations in the brain are linked to the quality of maternal attachment to the child. As the experience of baby blues was assessed 3 weeks after childbirth, it is reasonable to assume that some of the participants no longer had the symptoms at the time of the survey, although they might have had them at the time of the MRI measurement. These limitations notwithstanding, the study involved a large sample of postpartum women (as part of the ongoing longitudinal study related to early recognition of PPD), controlling for psychiatric history or the development of PPD and postpartum adjustment disorder within the first 12 weeks postpartum.

### Conclusion and perspectives

Based on our results and those of previous studies, we conclude that the exact biological/physiological function of the changes in brain morphology observed in humans shortly after childbirth remains unclear. Although, given that these changes have been seen to occur both in humans and other mammals, it can be safely assumed that they have a role to play. A profound impact of motherhood on the brain organization is already apparent in gestating mice (4). Animal studies have shown that increased interactions with pups over time during the early postpartum period lead to additional structural changes in the maternal brain (19, 50, 51). It is reasonable to assume that similar structural changes occur in the brain of human mothers. In view of the fact that the acute pregnancy-related GMV differences were not found to predict the MPAS scores, it would be interesting to investigate the adaptive neuroplasticity of the mother’s brain with respect to the quality of maternal attachment to the child over a longer postpartum period. In addition, evidence suggests that the cognitive functions may also be influenced by pregnancy (21). However, it is unclear if the reduction of brain volume in pregnancy is necessarily related to the deterioration of cognitive or other functions. On the contrary, decreased GMV may result in more communication both between and within brain regions (21). Another intriguing but as yet unanswered question is whether pregnancy (multiple pregnancies in particular) can have lifelong consequences with respect to brain health and the aging process. For instance, parity-induced endocrine changes have been suggested to influence brain response to sex hormones later in life (21), leaving a long-lasting footprint on the endocrine system. Large-scale epidemiological studies have reported associations between increasing parity and the risk of metabolic syndrome (52–54) and type 2 diabetes mellitus (55, 56). In addition, the physiological adaptations associated with pregnancy can render pregnant women susceptible not only to psychiatric but also, and more frequently, to metabolic and cardiovascular diseases. Conditions such as pre-eclampsia, pregnancy-induced hypertension and gestational diabetes are characterized by physiological responses indicative of the metabolic syndrome, potentially heralding future cardiovascular and metabolic diseases. However, the extent to which preexisting or pregnancy-induced medical conditions, multiparity or the age of women at the time of pregnancy, interact with pronounced GMV reductions in the brain during pregnancy, and their restoration after childbirth, remains unclear. Moreover, physiological changes such as the drastic decline in hormones, blood loss, changes in blood volume and blood pressure during labor and delivery may have an additional influence on the morphological alterations in the maternal brain.

Longitudinal studies are needed to understand the way the networks and brain structures that undergo pregnancy-related changes recover after childbirth, and which factors influence the recovery processes. Future studies tracking ovarian hormones from before pregnancy through pregnancy and the postpartum period, as well as longitudinal multimodal neuroimaging, cognitive and lifestyle assessments, will likely help identify the factors that contribute to the observed neuroanatomical changes during pregnancy and the postpartum restoration of brain structure. In addition, further studies are needed to investigate if the changes associated with pregnancy contribute to the development of psychiatric conditions, especially in women at higher risks for postpartum psychiatric disorders (e.g. women with a psychiatric history). Finally, the cumulative effect of multiple pregnancies on brain anatomy (if proven) may suggest that pregnancy has longterm consequences with respect to the maternal brain. And if pregnancy is indeed found to have negative long-term effects on aging, mental health or any other aspect of women’s health, it would be imperative to explore how these effects can be counterbalanced by such factors as healthy life style, cognitive activities, and, importantly, the prevention or early treatment of pregnancy-related somatic and psychiatric disorders.

## Materials and Methods

### Participants included in the analysis

The present study included a group of 78 non-depressed postpartum women (age range 21 to 42 years). The imaging data of these participants were selected from the brain imaging data pool (N = 174) of an ongoing longitudinal study pertaining to the early detection of postpartum depression (“Risk of Postpartum Depression-Study”, RiPoD) based on the following criteria: Edinburgh Postnatal Depression Scale (EPDS) scores below 10 immediately after childbirth (57) and no signs of postpartum psychiatric disorder based on the clinical interview at 12 weeks postpartum. The postpartum women included in the current analysis were further split into two groups based on their parity: 41 primiparous women (age range 22 to 39 years) and 37 multiparous women (age range 21 to 42 years). Within the multiparous group, 33 women had 2 children and 4 women had 3 children. The structural MRI session took place between one and four days after childbirth (M = 2.58 days postpartum, SD = 0.79 days postpartum).

### RiPoD Study

All participants of the RiPoD study were recruited in the Department of Gynecology and Obstetrics at the University Hospital Aachen within one to six days of childbirth. Twelve weeks after childbirth, the participants were invited to a final semi-standardized clinical interview for the final diagnosis by an experienced psychiatrist or psychologist. Women with current depression at the moment of recruitment, abuse of alcohol, drugs, psychotropic substances, antidepressant or antipsychotic medication during pregnancy, history of psychosis or manic episodes were excluded from the RiPoD study. In addition, only mothers of healthy children (determined by the routine German Child Health tests (U2) conducted within the first 3 to 10 days of life) were included. Barring the pregnancy- and child-related exclusion criteria, the recruitment was equitable and inclusive, representing a cross section of the local population. As part of the RiPoD study (for a comprehensive study overview please refer to 58), all participants had to complete the Edinburgh Postnatal Depression Scale (23), a 10-item self-report to assess depressive symptomatology in the postpartum period, immediately after childbirth and every three weeks for 12 weeks postpartum. Using the MPAS (22), a total score of maternal attachment including sub-scores for quality of attachment, absence of hostility and pleasure in interaction, was evaluated at 3, 6, 9 and 12 weeks postpartum. At 3 weeks postpartum, the experience of baby blues symptoms was assessed with both a self-report of experiencing baby blues and a cut-off score of > 10 on the Maternity Blues Questionnaire (59, 60) (see Figure 4).

**Figure 4.**
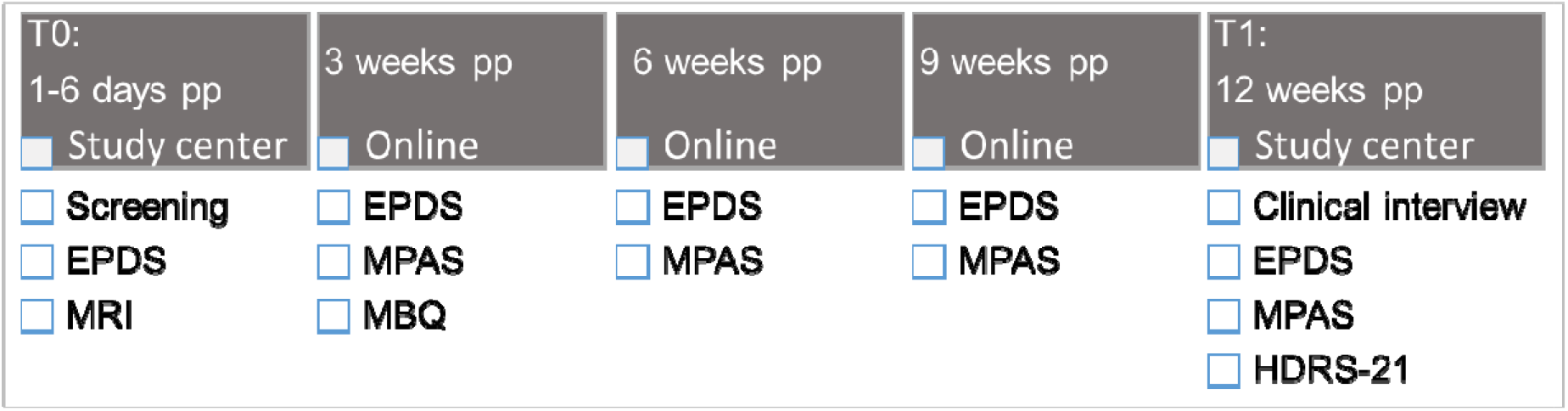
Excerpt from the RiPoD study procedure used for this study. Pp: postpartum; EPDS: Edinburgh Postnatal Attachment Scale, MRI: Magnetic resonance imaging; MPAS: Maternal Postnatal Attachment Scale; MBQ: Maternity Blues Questionnaire; HDRS-21: Hamilton Depression Scale.

### Nulliparous control subjects

The nulliparous control subjects were 40 healthy female adults (age range 19 to 35 years, M = 26.7 years, SD = 4.53 years) recruited through advertisement in the study center. Control subjects had no history of psychiatric disorders (identified by the short version of the German Structured Clinical Interview for DSM 4 Disorders (SCID-I) (61)) and had never been pregnant before, regardless of whether there was a (spontaneous) abortion or pregnancy outcome (e.g. extrauterine pregnancy).

### Behavioral data analysis

The analysis was conducted using SPSS® 25 IBM Corporation, Armonk NY, USA) for Windows®. The postpartum groups (primiparous and multiparous) were compared with regard to age, gestational age, child’s birth weight, birth mode, EPDS scores and MPAS total as well as subscale scores. For each of the continuous measures, independent sample t-tests were conducted. In case of violation of the assumption of homogeneity of variance (Levene’s test), t-statistic not assuming equality of variance was computed. For categorical measures, chi square tests were conducted.

Separate ANOVAs for repeated measures were calculated within each parity group with measurement time points after birth and EPDS scores or MPAS total scores and subscale scores as within-subject variables. The Greenhouse-Geisser correction was used to adjust degrees of freedom when significant non-sphericity was detected via the Mauchly’s test. The significant findings were pursued with Bonferroni-corrected pairwise comparisons. The effect sizes of the significant results are reported using partial eta squared (η_p_^2^) for F-tests (small: 0.20-0.05, medium: 0.06−.013, large: .14 and greater) and Cohen’s d for pairwise comparisons (small: 0.20–0.49, medium: 0.50–0.79, large: 0.80 and greater) (62).

A significance level of p-value less than .05 was used.

### MRI data acquisition

Neuroimaging data were acquired using a 3 Tesla Prisma MR Scanner (Siemens Medical Systems, Erlangen, Germany) located in the Medical Faculty of RWTH Aachen University. T1-weighted structural images were acquired by means of a three-dimensional magnetization-prepared rapid acquisition gradient echo imaging (MPRAGE) sequence (4.12 minutes; 176 slices, TR = 2300 ms, TE = 1.99 ms, TI = 900 ms, FoV = 256 × 256 mm^2^, flip angle = 9°, voxel resolution = 1 × 1 × 1 mm^3^).

### Voxel-based morphometry (VBM)

Anatomical imaging data were preprocessed using the Computational Anatomy Toolbox (CAT12 Version r1728) and SPM12 toolbox implemented in Matlab 2015b (MathWorks, Inc., Natick, MA). The default settings of CAT12 were applied for spatial registration, segmentation and normalization (63).

All images were affine registered to standard tissue probability maps (TPM) by correcting individual head positions and orientations, and translated into Montreal Neurologic Institute (MNI) space. The images were segmented into gray matter, white matter, and cerebrospinal fluid. For normalization, the Diffeomorphic Anatomic Registration Through Exponentiated Linear algebra algorithm (DARTEL) was used (64) as DARTEL affords a more precise spatial normalization to the template than the conventional algorithm (65). A homogeneity check identified no outliers, thus the gray matter volumes (GMV) of all participants were included in subsequent analyses. Finally, the modulated GMV were smoothed with an 8 mm FWHM Gaussian kernel.

### Surface-based morphometry (SBM)

The CAT12 toolbox was used to extract cortical thickness information. Volumes were segmented using surface and thickness estimation in the writing options. Local maxima were projected to the gray matter voxels by using neighbor relationship described by the WM distance, equaling cortical thickness. The estimation of cortical thickness was performed based on projection-based thickness including partial volume correction, sulcal blurring and sulcal asymmetries without sulcus reconstruction (66). Topological correction was performed through an approach based on spherical harmonics. For inter-participant analysis, an algorithm for spherical mapping of the cortical surface was included (67). An adapted volume-based diffeomorphic DARTEL algorithm was then applied to the surface for spherical registration. All scans were re-sampled and smoothed with a Gaussian kernel of 15 mm FHWM.

### Statistical analyses for VBM and SBM

The statistical analysis was performed using SPM 12. Smoothed gray matter segments (VBM analysis) of all participants were implemented in a voxel-wise whole-brain one-way ANOVA. To test whether the baby’s gender or the experience of baby blues (yes/no) had an effect on GMV, whole-brain full-factorial general linear model (GLM) with factor group (two levels) and factor baby blues (2 levels) or gender (2 levels) was performed. The effect of hormonal contraception intake (yes/no) in nulliparous control subjects was pursued with a two-sample t-test. In all VBM analysis, age and total intracranial volume (TIV) were used as control variables. Gray matter structures in VBM analysis are labeled with reference to the Anatomy Toolbox for SPM (68) and the Automated anatomical labelling atlas 3 (69). For the SBM analysis, resampled and smoothed thickness files of all participants were inserted in a whole-brain one-way ANOVA. Here, only age was used as a control variable. Structures are labeled with reference to the Desikan-Killiany atlas (70).

The statistical threshold was set at p < .05 cluster-level FWE-correction, with a cluster-forming threshold at voxel-level p < .001, k = 920 voxels for VBM and k = 116 voxels for SBM. All results are presented in the MNI space.

### Multivariate regression analyses

The PRONTO toolbox 2.0 (http://www.mlnl.cs.ucl.ac.uk/pronto/) implemented in Matlab was used to predict the MPAS total and its attachment, hostility and interaction subscales using whole-brain voxel-wise GMV and controlling for age and TIV. A kernel ridge regression was computed using a leave-one-out approach with the default settings. Permutation-based non-parametric p-values (1000 permutations) were computed for the correlation between observed and predicted scores.

## Acknowledgments

This study was funded by the rotation program (2015-2017) of the medical faculty of the University Hospital RWTH Aachen, the Deutsche Forschungsgemeinschaft (DFG, No. 410314797).

This work was supported by the Brain Imaging Facility of the Interdisciplinary Center for Clinical Research (IZKF) Aachen within the Faculty of Medicine at RWTH Aachen University.

## Supplementary Information

**SI Whole-brain VBM results**

**Table S1.** Brain regions showing GMV differences in control subjects > primiparous women (t-contrast from random-effects GLM, p < .05, cluster-level FWE correction, k = 920)

| Anatomical region | Peak BA | Side | k | Peak voxel |  |  |  |
| --- | --- | --- | --- | --- | --- | --- | --- |
|  |  |  |  | T | x | y | z |
| Cerebellum vermis VIII, VI, IX, Crus 1, fusiform gyrus, inferior occipital gyrus, inferior temporal gyrus, lingual gyrus, inferior occipital gyrus, calcarine gyrus, middle temporal gyrus, parahippocampal gyrus | 18, 19, 20 37 | R/L | 20607 | 5.56<br>5.31<br>5.28 | 6<br>-5<br>-20 | -59<br>-60<br>-71 | -30<br>-30<br>-23 |
| Pallidum, caudate, rectal gyrus, thalamus, cerebellum Crus 2, vermis VII, X, lingual gyrus, inferior occipital gyrus, calcarine gyrus, middle temporal gyrus, parahippocampal gyrus, anterior cingulate cortex, putamen, superior medial frontal gyrus, olfactory sulcus, medial orbital gyrus, superior temporal gyrus, middle frontal gyrus, amygdala, insula, midcingulate cortex, fusiform gyrus, hippocampus, inferior frontal gyrus (p. triangularis and p. orbitalis) | 11, 25, 23, 34 | R/L | 12159 | 5.17<br>4.94<br>4.80 | 12<br>20<br>-12 | 3<br>17<br>3 | -3<br>-12<br>-6 |
| Superior parietal lobule, precuneus midcingulate cortex, postcentral gyrus, paracentral lobule, cuneus | 6, 7, 24, 31 | R/L | 3988 | 5.84<br>5.57<br>4.68 | -26<br>-15<br>-8 | -60<br>-60<br>-38 | 50<br>50<br>45 |

Note. MNI coordinates and T-values of first three peak voxels of the significant clusters and a maximum of 4 Brodmann areas (BA) within the clusters are shown. Anatomical regions are in the order of the peaks within the cluster. L = left hemisphere, R = right hemisphere.

**Table S2.** Brain regions showing GMV differences in control subjects > multiparous women (t-contrast from random-effects GLM, p < .05, cluster-level FWE correction, k = 920).

| Anatomical region | Peak BA | Side | k | Peak voxel |  |  |  |
| --- | --- | --- | --- | --- | --- | --- | --- |
|  |  |  |  | T | x | y | z |
| Cerebellum vermis VI, VIII, Crus 1, fusiform gyrus, Heschl's gyrus, superior and middle temporal gyrus, lingual gyrus, precentral gyrus, rolandic operculum, inferior temporal gyrus, parahippocampal gyrus, hippocampus, inferior frontal gyrus (p. opercularis), insula, inferior occipital gyrus, postcentral gyrus, calcarine, superior temporal pole | 18, 19, 22, 37 | R/L | 36157 | 6.45<br>6.37<br>5.81 | -17<br>27<br>-20 | -71<br>-62<br>-50 | -24<br>-27<br>-12 |
| Medial temporal pole, thalamus, putamen, middle temporal gyrus, amygdala, pallidum, superior and inferior temporal gyrus, caudate, olfactory sulcus, insula, rolandic operculum, rectal gyrus, parahippocampal gyrus, Heschl's gyrus, medial orbital gyrus, hippocampus, inferior frontal gyrus (o. orbitalis) precentral gyrus | 21, 25, 34, 38 | R/L | 7842 | 5.10<br>4.94<br>4.82 | 39<br>-2<br>48 | 6<br>-12<br>11 | -33<br>-2<br>-26 |
| Middle frontal gyrus, superior medial frontal gyrus, middle orbital gyrus, anterior cingulate gyrus | 9, 10, 32 | R/L | 4838 | 5.14<br>5.03<br>4.97 | -20<br>14<br>11 | 47<br>50<br>56 | 29<br>18<br>21 |
| Superior parietal lobule, middle occipital gyrus, postcentral gyrus, inferior parietal lobule, precuneus, postcentral gyrus, supramarginal gyrus | 7, 19, 40 | L | 4300 | 6.34<br>5.71<br>5.53 | -24<br>-17<br>-15 | -60<br>-63<br>-62 | 47<br>42<br>48 |
| Rolandic operculum, inferior frontal gyrus (p. triangularis, p. opercularis, p. orbitalis), insula, middle frontal gyrus, superior temporal pole, precentral gyrus, Heschl's gyrus, putamen | 9, 13, 22, 44 | R | 3520 | 5.37<br>5.08<br>4.94 | 44<br>54<br>48 | 9<br>8<br>5 | 12<br>0<br>14 |
| Supplementary motor cortex, superior medial frontal gyrus, midcingulate cortex | 6, 8, 32 | R/L | 1972 | 5.05<br>4.79<br>4.75 | 0<br>5<br>3 | 5<br>14<br>17 | 60<br>68<br>62 |
| Middle and superior occipital gyrus, angular gyrus, superior parietal gyrus | 7, 19, 39 | R | 1379 | 4.00<br>3.98 | 30<br>30 | -83<br>-81 | 20<br>24 |

Note. MNI coordinates and T-values of first three peak voxels of the significant clusters and a maximum of 4 Brodmann areas (BA) within the clusters are shown. Anatomical regions are in the order of the peaks within the cluster. L = left hemisphere, R = right hemisphere.

**Table S3.**
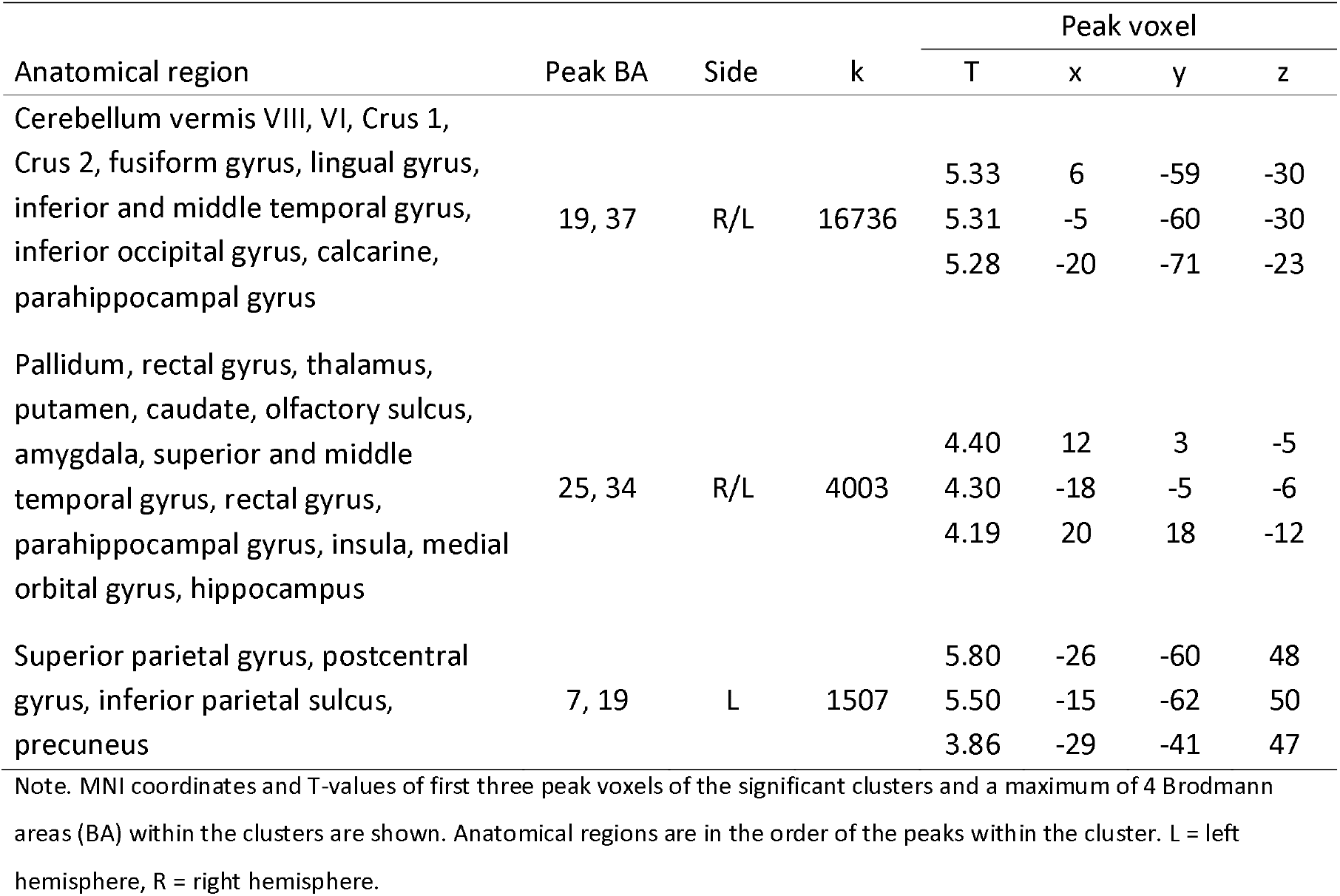
Brain regions showing GMV differences in the conjunction analysis (controls > primiparous ⍰ controls > multiparous, p < .05, cluster-level FWE correction, k = 920).

| Anatomical region | Peak BA | Side | k | Peak voxel |  |  |  |
| --- | --- | --- | --- | --- | --- | --- | --- |
|  |  |  |  | T | x | y | z |
| Cerebellum vermis VIII, VI, Crus 1,<br>Crus 2, fusiform gyrus, lingual gyrus,<br>inferior and middle temporal gyrus,<br>inferior occipital gyrus, calcarine,<br>parahippocampal gyrus | 19, 37 | R/L | 16736 | 5.33<br>5.31<br>5.28 | 6<br>-5<br>-20 | -59<br>-60<br>-71 | -30<br>-30<br>-23 |
| Pallidum, rectal gyrus, thalamus,<br>putamen, caudate, olfactory sulcus,<br>amygdala, superior and middle<br>temporal gyrus, rectal gyrus,<br>parahippocampal gyrus, insula, medial<br>orbital gyrus, hippocampus | 25, 34 | R/L | 4003 | 4.40<br>4.30<br>4.19 | 12<br>-18<br>20 | 3<br>-5<br>18 | -5<br>-6<br>-12 |
| Superior parietal gyrus, postcentral<br>gyrus, inferior parietal sulcus,<br>precuneus | 7, 19 | L | 1507 | 5.80<br>5.50<br>3.86 | -26<br>-15<br>-29 | -60<br>-62<br>-41 | 48<br>50<br>47 |

Note. MNI coordinates and T-values of first three peak voxels of the significant clusters and a maximum of 4 Brodmann areas (BA) within the clusters are shown. Anatomical regions are in the order of the peaks within the cluster. L = left hemisphere, R = right hemisphere.

**Results of whole-brain SBM**

**Table S4.**
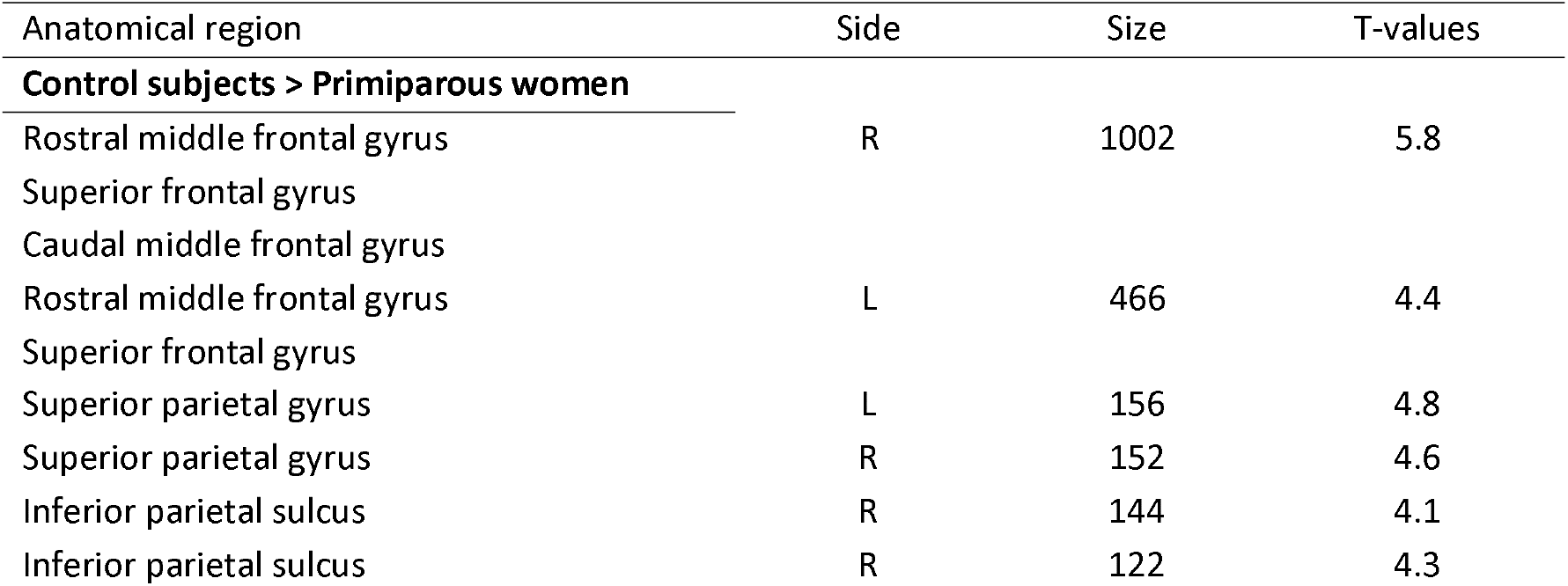
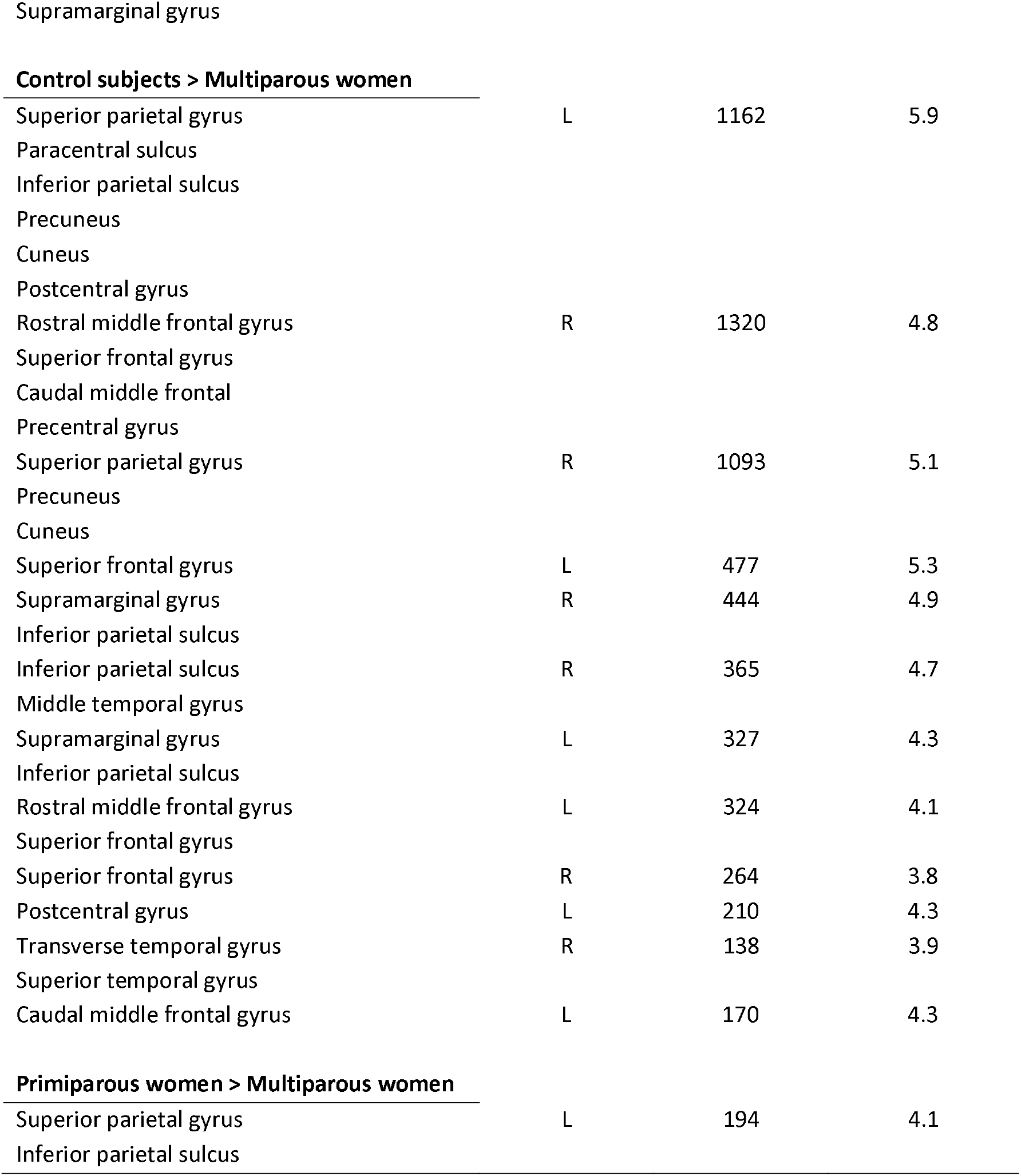
Overview of cortical thickness differences. Atlas labeling was performed according to the Desikan-Killiany atlas (37)

